# Tracing the intruders: a global appraisal of marine invasive species detection through DNA-based approaches

**DOI:** 10.64898/2026.05.05.722998

**Authors:** Sofia Duarte, Filipe O. Costa

## Abstract

Early detection and monitoring of non-indigenous species (NIS) is crucial to prevent their establishment and to reduce ecological and economic impacts in coastal ecosystems. Traditional monitoring approaches, which rely largely on morphological identification of collected organisms, are often time-consuming and may fail to detect species that occur at low abundance, are morphologically cryptic, or are present in the form of inconspicuous life stages. DNA-based approaches, particularly those resorting to environmental DNA, have demonstrated high aptitude for biodiversity monitoring and biosecurity surveillance. By examining the genetic material from bulk community samples or released into the environment, DNA-based approaches enable the detection of species without the need for direct observation, thereby increasing detection sensitivity and expanding the scope of monitoring programs. Despite the rapid growth of its employment in marine monitoring, a global synthesis of the status and trends of DNA-based approaches for detecting NIS in this environment has been lacking. Here, we present such synthesis, based on 146 published studies employing DNA for NIS detections in coastal environments. Two main methodological approaches were used across the reviewed studies, namely DNA metabarcoding which was applied in 49% of studies, closely followed by targeted single-species PCR assays, used in 42% of the studies. A smaller proportion of studies (10%) combined both approaches, integrating broad community screening with targeted detection to improve surveillance efficiency. Globally, 752 NIS were detected across disparate taxonomic groups, with metazoans representing the largest proportion of detections (464 species), followed by Chromista (210 species) and Plantae (77 species). Among these, the most frequently detected taxonomic groups included Dinophyceae (Dinoflagellata), Teleostei (Chordata), Florideophyceae (Rodophyta), Polychaeta (Annelida), Copepoda and Malacostraca (Arthropoda), and Ascidiacea (Chordata). At the species level, several well-known marine invaders were recurrently reported, including *Bugula neritina* (Linnaeus, 1758), *Styela plicata* (Lesueur, 1823), *Acartia (Acanthacartia) tonsa* Dana, 1849-1852, and *Botryllus schlosseri* (Pallas, 1766), highlighting the ability of DNA approaches to detect widespread and established invaders across different regions. The mitochondrial cytochrome c oxidase subunit I (COI) gene was the most widely used genetic marker, reflecting its broad taxonomic coverage and extensive representation in reference databases, particularly for targeting Metazoa. Ribosomal RNA genes, particularly 18S and 16S rRNA gene markers, were also frequently employed to target a wider range of eukaryotic taxa. Regarding sampled substrates, water was by far the most analyzed substrate, followed by zooplankton and biofouling communities collected from man-made structures. Notably, approximately 31% of all NIS detections reported in the reviewed studies constituted new regional records. These results highlight the potential of eDNA for coastal monitoring but also underline important limitations. Persistent geographical, taxonomic, and methodological biases can affect detection outcomes, and reliance on single sample types or markers may increase false negatives - particularly critical for NIS early detection. Therefore, multi-marker and multi-substrate approaches are essential to improve detection reliability and support effective biosecurity strategies. As reference databases continue to expand and methodological protocols become increasingly standardized, DNA-based monitoring is likely to play a central role in future management and surveillance of biological invasions in coastal ecosystems.

**Graphical Abstract:** 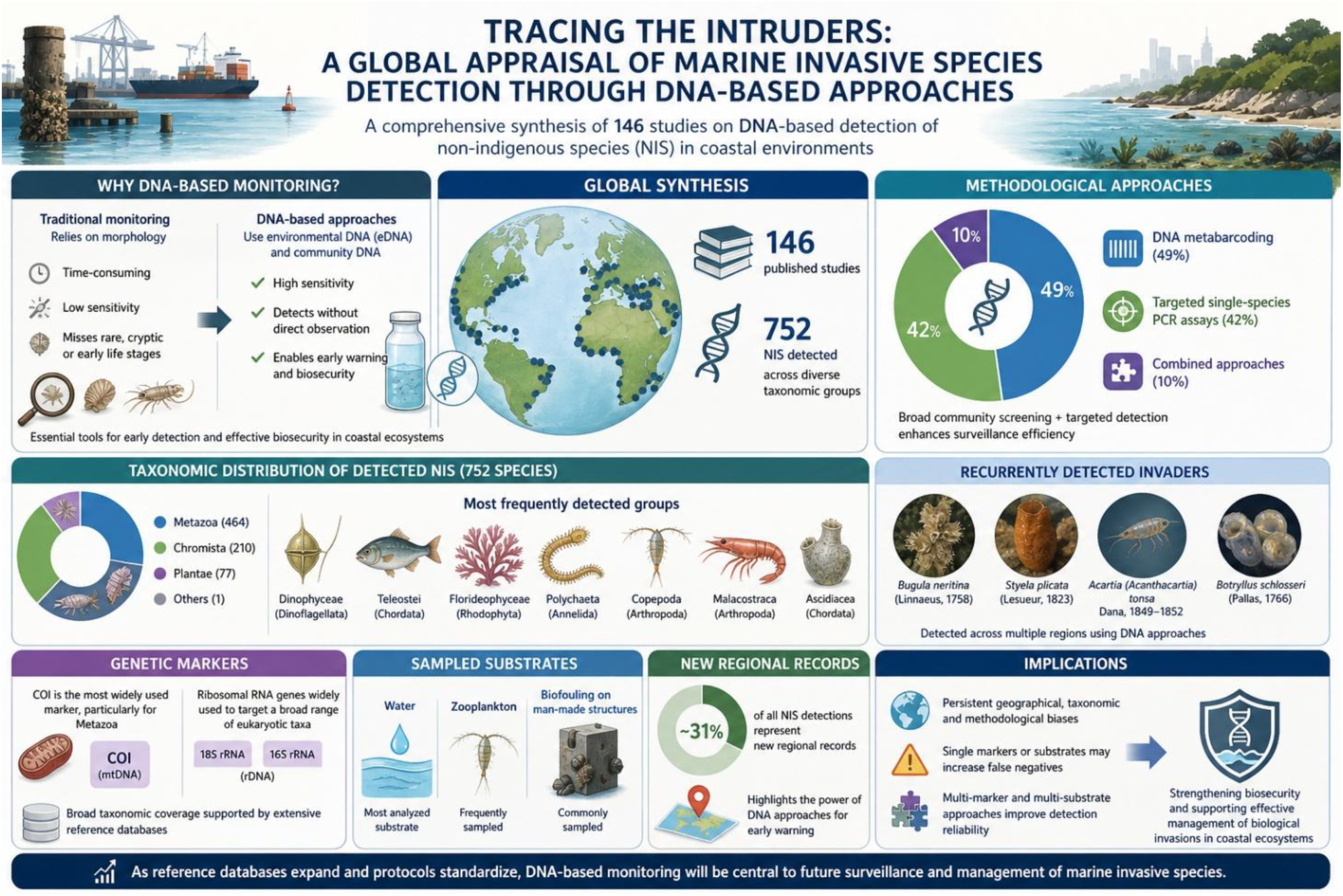

## 1. Introduction

Biological invasions are among the leading drivers of biodiversity loss and ecosystem change in marine and coastal environments worldwide, with profound ecological, economic, and socio-economic consequences (Carlton, 1996; Molnar et al., 2008; Seebens et al., 2017). Coastal ecosystems are particularly vulnerable to non-indigenous species (NIS) introductions due to intense human activity, including global shipping, aquaculture, recreational boating, coastal development, and plastic pollution, which can act as a vector for rafting organisms, collectively increasing propagule pressure and facilitating secondary spread (Audrézet et al., 2021; Ruiz et al., 2000; Seebens et al., 2013). Once established, marine NIS can alter food webs, displace native species, modify habitats, and impair ecosystem services, often with impacts that are difficult or impossible to reverse (Grosholz, 2002; Katsanevakis et al., 2014).

Early detection remains one of the most effective strategies to prevent the establishment and spread of invasive species and to reduce the costs and ecological damage associated with late-stage management (Lodge et al., 2016; Simberloff et al., 2013). However, traditional monitoring approaches based on morphology, visual surveys, and trapping often fail to detect invaders at low abundance, early life stages, or when species are cryptic or morphologically indistinguishable from native taxa (Audrézet et al., 2021; Darling and Blum, 2007; Zaiko et al., 2015). These limitations are particularly acute in marine systems, where high biodiversity, taxonomic expertise gaps, and logistical constraints hinder comprehensive surveillance.

Over the past decade, DNA-based tools approaches which enables the identification of species within a sample by sequencing short, taxonomically informative DNA regions, have emerged as powerful alternatives or complements to conventional monitoring methods (Deiner et al., 2017; Ficetola et al., 2008; Ruppert et al., 2019). This has proven especially valuable for identifying rare, elusive, or early-stage invaders, often with higher sensitivity than traditional surveys (Darling, 2019; Duarte et al., 2023a, 2021; Goldberg et al., 2016). DNA-based monitoring has therefore become increasingly integrated into marine biosecurity frameworks and management strategies worldwide (Darling and Mahon, 2011; Wood et al., 2013).

Two main molecular methodologies dominate DNA-based NIS detection. Species-specific assays, typically based on qPCR or droplet digital PCR (ddPCR), target and amplify DNA from a single species using highly specific primers and probes, allowing for precise and highly sensitive detection and quantification of priority taxa. These approaches are particularly suited for early warning and targeted surveillance (Guri et al., 2024; Takahara et al., 2012; Wood et al., 2019a). In contrast, DNA metabarcoding relies on the amplification and high-throughput sequencing of standardized genetic markers from mixed DNA samples using universal primers, enabling the simultaneous detection of multiple taxa. This provides a community-wide snapshot that can reveal unexpected or previously unrecorded NIS alongside native biodiversity (Lavrador et al., 2024; Simmons et al., 2016; Zaiko et al., 2015). While each approach has distinct advantages and limitations, their complementary use has the potential to substantially enhance the effectiveness of marine invasion monitoring. These approaches can be applied both to bulk samples (e.g., macrozoobenthos) and to environmental DNA (eDNA), defined as the genetic material released into the environment through sloughed cells, gametes, feces, or decomposing tissues, allowing non-invasive detection of organisms without direct observation or capture (Kavakiotis et al., 2026).

Despite the rapid expansion of eDNA applications in marine invasion science, existing syntheses have largely focused on methodological performance, case studies, or specific environments, often with a regional or taxonomic bias (Belle et al., 2019; Coble et al., 2019; Duarte et al., 2023a; Schenekar, 2023). To date, there has been no global, taxonomically inclusive synthesis quantifying which non-indigenous species have been detected using DNA in marine and coastal ecosystems, nor how methodological choices (e.g. markers, substrates, platforms) shape detection outcomes. Such a synthesis is essential to assess the real-world contribution of DNA-based tools to early detection, to identify geographic and taxonomic gaps, and to inform the design of future monitoring programs.

Here, we present a global synthesis of non-indigenous species detected using DNA-based approaches in marine and coastal ecosystems, based on 146 peer-reviewed studies published between 2010 and 2024. Specifically, we (i) quantify the number and taxonomic composition of NIS detected using DNA-based tools, (ii) examine geographic patterns in research effort and detection outcomes, (iii) evaluate methodological trends in detection strategies, sample types, and genetic markers, and (iv) assess the contribution of DNA to the discovery of new regional NIS records. By integrating results across taxa, regions, and methodological approaches, this review provides a comprehensive benchmark for the current state of DNA-based NIS surveillance and offers guidance for the development of more effective and harmonised monitoring strategies in marine and coastal environments.

## 2. Materials and Methods

### 2.1. Literature search and dataset selection

A comprehensive literature search was performed in the Web of Science database on January 2nd, 2025, to compile a state-of-the-art overview of eDNA-based tools for the detection and monitoring of non-indigenous species (NIS) in marine and coastal ecosystems. The search was restricted to titles, abstracts, and keywords, using combinations of the following terms:1) eDNA-related terms: “metabarcoding”, “high throughput sequencing”, “next generation sequencing”, “polymerase chain reaction”, “PCR”, “environmental DNA”, “eDNA”; 2) NIS-related terms: “invasive species”, “non indigenous species”, “alien species”, “exotic species”, “non native species”, “marine pest”, “nuisance species” and 3) Ecosystem-related terms: “coast”, “marine”, “estuar*”, “brackish”, “transition*”, “sea*”, “lagoon”, “ballast water”, “biofouling”*. To focus on experimental work, the “exclude review articles” option was applied. The search yielded 408 articles published between 2010 and 2024 (**Table S1**). An additional 21 relevant articles from personal collections (published between 2015 and 2022) were also included, as they were not retrieved through the search (**Table S1**).

Following individual screening, 31 records were excluded for being review articles (despite the exclusion filter) or not involving DNA-based approaches. The remaining 394 studies were assessed for eligibility since they employed DNA-based tools. Of these, 248 were excluded for one or more of the following reasons: 1) they focused on general biodiversity assessments; 2) examined the diets of NIS; 3) addressed microbial communities (e.g., bacteria, viruses); 4) conducted population genetics analyses; 5) were gap-analyses and 6) were carried out in freshwater or terrestrial environments. A total of 146 studies met the inclusion criteria and were retained for further analysis (**Fig. 1, Table S2**).

**Fig. 1.**
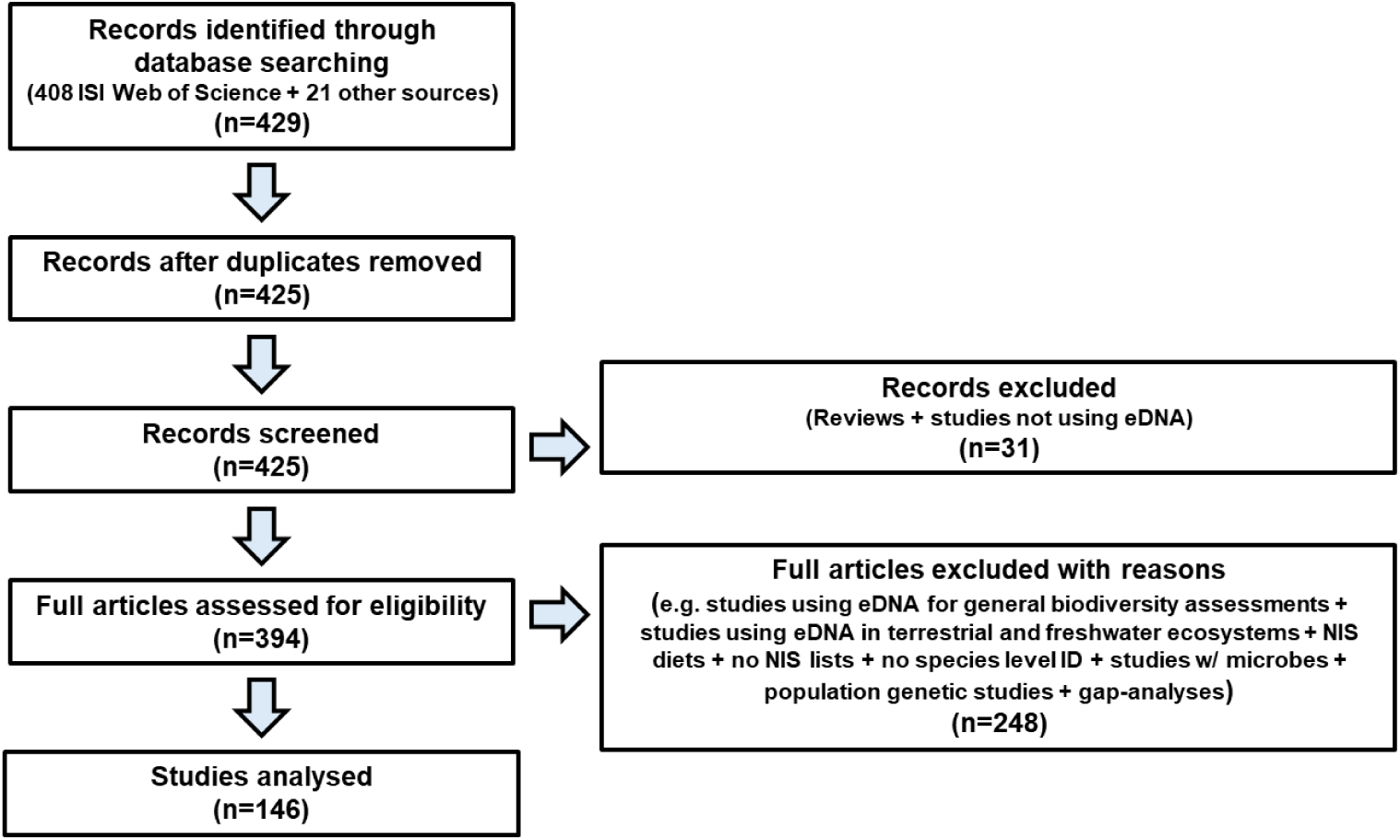
Diagram illustrating the selection process of publications included in the literature review on the use of DNA based methods for the detection and monitoring of non-indigenous species in marine and coastal ecosystems.

### 2.2. Data analysis

From each selected publication, the following information was extracted: 1) country/geographic region; 2) habitat (e.g., open sea, recreational marina, lagoon); 3) detection method (e.g., metabarcoding, qPCR, ddPCR); 4) sample type (e.g., water, sediment, zooplankton, fouling communities); 5) genetic marker (and length in bp); 6) primers and probes (in the case of qPCR or dPCR); 7) non-indigenous species detected (for this results sections, supplementary material, tables were fully inspected in order to get as much completed as possible) and 8) other information that we considered relevant for the study.

All information was compiled and standardized in Microsoft Excel (Supplementary material, **Table S2**). Descriptive analyses were conducted using dynamic pivot tables to summarize patterns in publication effort, geographic coverage, methodological approaches, genetic markers, sampled substrates, and taxonomic composition of detected non-indigenous species (NIS). Temporal trends in publication output were assessed by quantifying the number of studies published per year and the cumulative number of publications over time. From 2017 onwards, a linear regression was fitted to cumulative publication counts to estimate the rate of increase in DNA-based NIS studies (GraphPad Prism 8.0.1).

The number of studies and NIS detections was quantified per region, taxonomic group, molecular approach (e.g. metabarcoding vs. single-species detection), genetic marker, and sample type. Taxonomy and nomenclature for Metazoa were checked and standardized using the World Register of Marine Species (WoRMS; https://www.marinespecies.org/, accessed on 30^th^ January 2026), which also allowed merging records for species reported under different synonyms or outdated names. For Chromista and Plantae (which were mostly algae), taxonomy and nomenclature were verified and standardized using AlgaeBase (https://www.algaebase.org/, accessed on 29^th^ April 2026).

Graphs illustrating spatial patterns, methodological trends, and taxonomic distributions were generated using GraphPad Prism 8.0.1. These included bar charts and proportional representations to visualize the relative contribution of regions, markers, substrates, and organismal groups to overall NIS detections. Venn diagrams (https://www.interactivenn.net/) were used to assess overlap in NIS detections among different sampled substrates (e.g. water, plankton, biofouling) and genetic markers, allowing identification of shared and substrate- or marker-specific detections. The proportion of NIS detections corresponding to new regional records was calculated based on information reported in the original studies.

## 3. Results and Discussion

### 3.1. Frontiers in Marine Science leads DNA-based NIS publications

Despite our efforts to conduct an exhaustive literature search, it is possible that a few relevant studies may have escaped our attention. Nevertheless, the inspection of 146 publications provided a robust dataset on the number of non-indigenous species detected to date using DNA-based tools in coastal and marine ecosystems. It also enabled the identification of major methodological trends, including the prevalence of active *versus* passive surveillance, the most used genetic markers and primers, and the detection platforms employed. A detailed analysis of the 146 selected articles revealed that research on the use of DNA-based tools for detecting non-indigenous species has been published across 57 scientific journals (**Table S2, Fig. S1**). However, only 28 of these journals appeared in more than 1% of the publications (i.e., at least two articles) (**Fig. S1**). The most frequently selected journals for publishing studies related to the use of DNA-based tools for detection and monitoring of marine NIS were Frontiers in Marine Science (11 publications), followed by Marine Pollution Bulletin and Scientific Reports (both with 9 publications) and Science of the Total Environment (8 publications) (**Table S2, Fig. S1**).

The earliest publication we identified on the use of eDNA-based tools for the detection of non-indigenous species (NIS) was by Willis and colleagues in 2011, who developed a PCR-based assay to enable the early detection of the sea squirt *Diplosoma listerianum* in Atlantic Canada (Willis et al., 2011). Since then, the number of publications in this field has steadily increased, with a particularly sharp rise from 2017 onwards, reaching an average rate of 17.2 publications per year (**Fig. 2a**). The first study to apply DNA metabarcoding for NIS detection was conducted by Zaiko and colleagues in 2015 who surveyed NIS in coastal areas of southeastern Baltic Sea (Zaiko et al., 2015). Although an earlier study by Pochon and colleagues (2013) evaluated the detection limits of next-generation sequencing for the surveillance and monitoring of international marine pests, it was not included in the current analysis because it was conducted using mock communities (Pochon et al., 2013). This highlights that one of the earliest applications of DNA-based tools in aquatic conservation was the detection of invasive species, driven by their high sensitivity and ability to enable early detection - both essential for timely management and control efforts (Ficetola et al., 2008; Pochon et al., 2013).

**Fig. 2.**
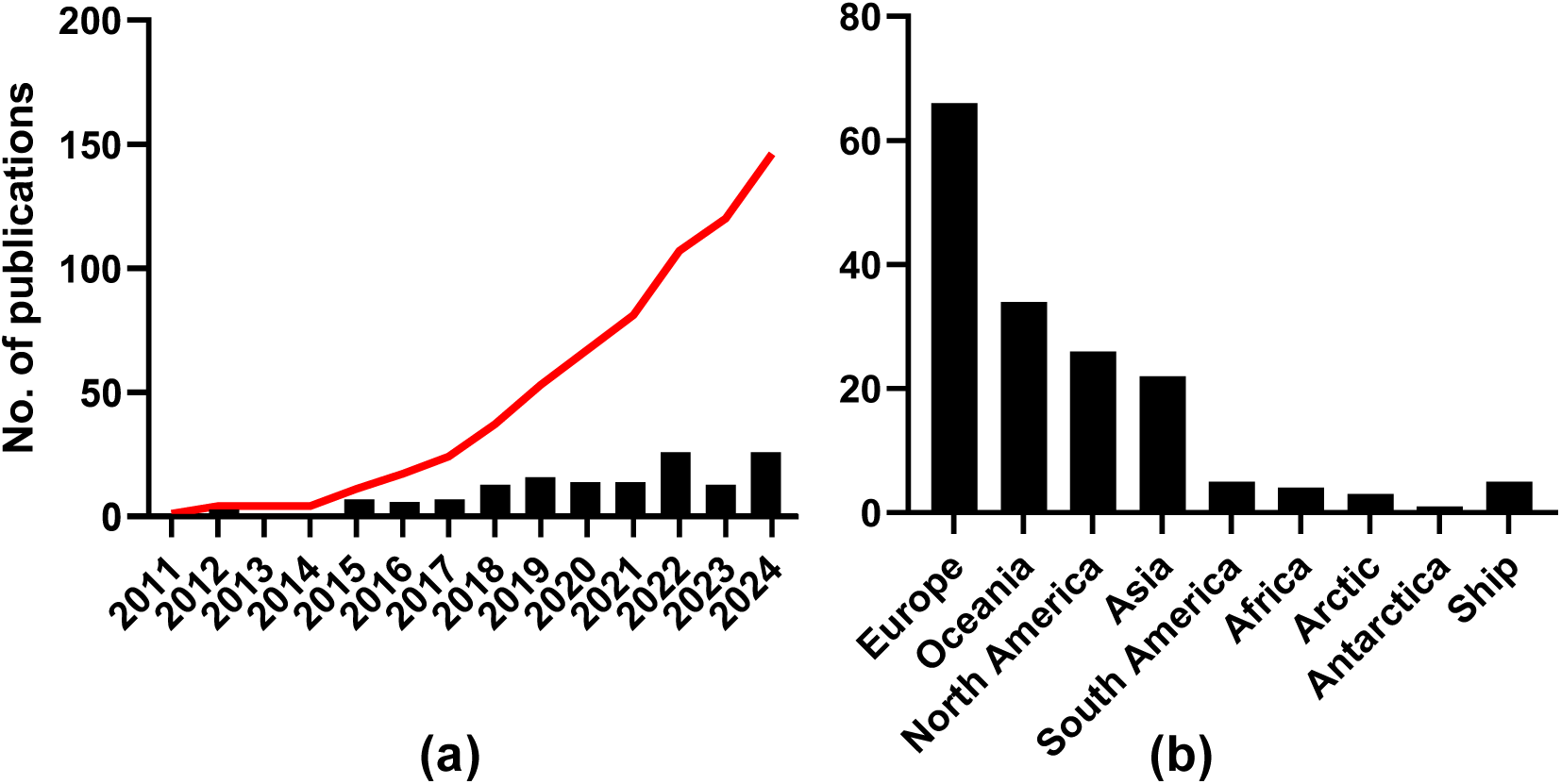
**(a)** Number of publications per year and cumulative number of publications (red line) and **(b)** number of publications per geographic regions employing DNA-based approaches for the detection and monitoring of non-indigenous species in marine and coastal ecosystems. From 2017 onwards the number of publications has increased at a rate of 17.2 papers per year (Linear regression, ±0.74, r^2^=0.9891).

### 3.2. DNA-based NIS monitoring has been applied mostly in Europe and Oceania

Most of the studies have been conducted in Europe (66 out of 146) (Couton et al., 2022; Holman et al., 2019; Suarez-Menendez et al., 2020), followed by Oceania (34) (Brand et al., 2022; Wood et al., 2019b; Zaiko et al., 2016), North America (26) (Grey et al., 2018; LeBlanc et al., 2020; Shang et al., 2019) and Asia (22) (Kim et al., 2020, 2018; Wang et al., 2022) (**Fig. 2b**). A geographic bias towards the Northern Hemisphere has been reported in several literature reviews regarding the adoption of DNA-based tools, particularly in freshwater environments (Belle et al., 2019; Coble et al., 2019; Schenekar, 2023). However, the current review suggests a different pattern in the context of invasive species detection, with a comparatively higher adoption in the Southern Hemisphere - especially in Oceania - where coastal ecosystems are heavily impacted by biological invasions. In fact, a closer look into the countries that were most surveyed revealed New Zealand in the top (20 studies), followed by Spain (16), Canada (14) and USA and Australia (both with 12) (**Fig. S2**). This is likely because these countries were among the first to adopt molecular methods to inform and support management decisions (Darling and Mahon, 2011; Xiong et al., 2016). New Zealand has one of the highest rates of non-indigenous species introductions in marine environments worldwide (Wood et al., 2013). Between 2009 and 2015, the cumulative number of marine non-indigenous species in the country increased by 10% (Inglis and Seaward, 2023). Similarly, in North America, the number of invasive species has more than tripled since the beginning of the 21st century (Leduc et al., 2019). The high number of studies in these countries may also reflect the presence of more active research communities working at the interface between molecular ecology and invasion science.

### 3.3. Metabarcoding and targeted-species detection have been applied with similar frequency in DNA studies

Among the 146 published studies, DNA metabarcoding was used in 49% of cases, single-species assays in 42%, while only 10% employed both approaches (**Table S2, Fig. 3**).

**Fig. 3.**
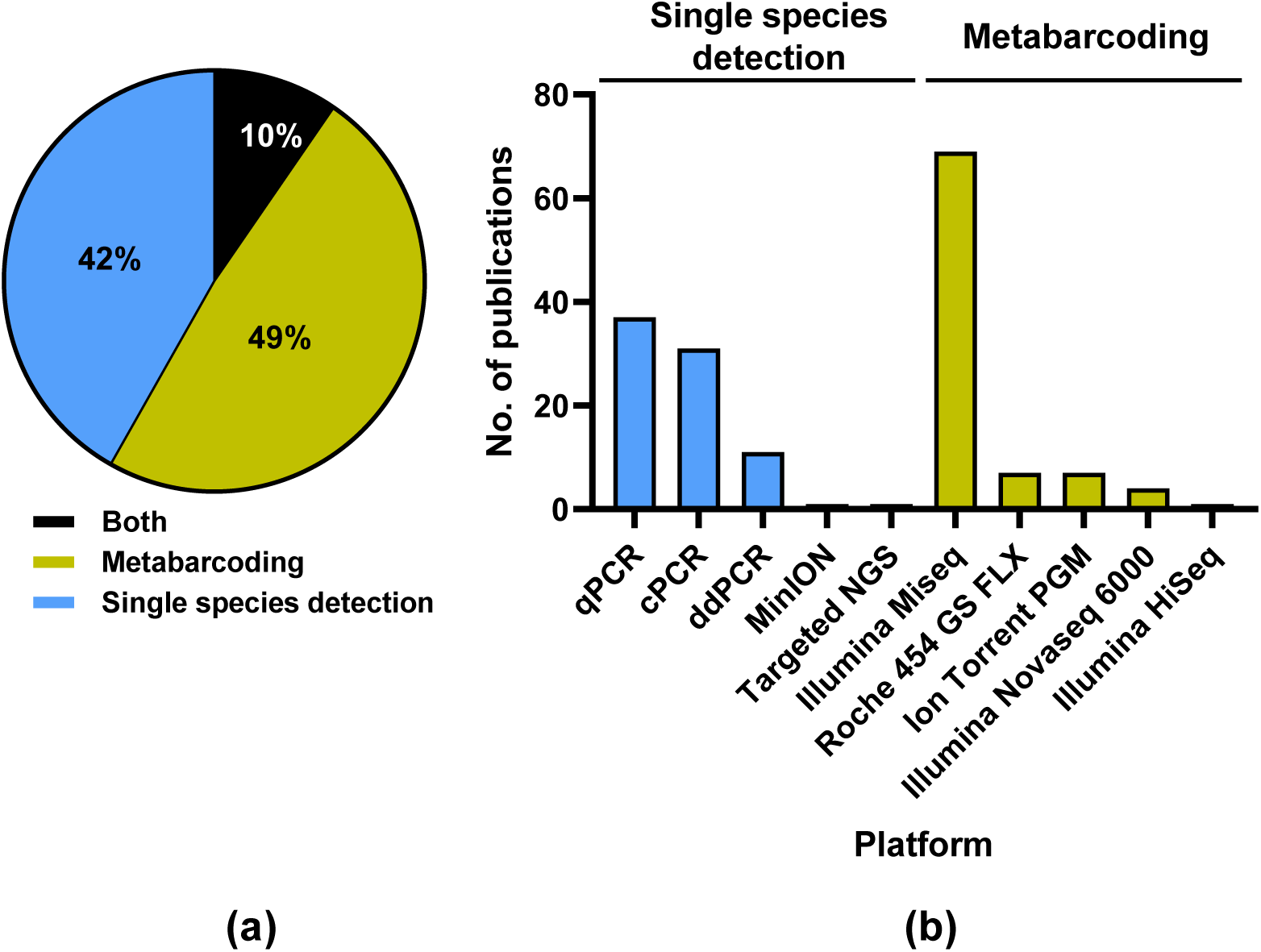
**(a)** % of publications using single-species detection, metabarcoding, or both strategies for non-indigenous species surveys in marine and coastal ecosystems, and **(b)** the platforms applied in each approach.

The choice between these molecular tools often depends on the specific objectives of the study and the stage of invasion being targeted. Single-species assays (e.g., qPCR, ddPCR) offer high sensitivity and specificity, making them particularly valuable for the early detection of priority invasive species, even at very low DNA concentrations (e.g. *Sabella spallanzanii* (Gmelin, 1791) in New Zealand coastal environments) (Brand et al., 2022; Wood et al., 2019a, 2019b, 2017). However, their main limitation lies in their taxonomic narrowness - each assay targets only one or a few species at a time, which restricts their use in broad biodiversity assessments or when the identity of potential invaders is unknown. In contrast, DNA metabarcoding provides a comprehensive, community-wide snapshot, enabling the simultaneous detection of multiple taxa, including unexpected or previously unrecorded non-indigenous species (Simmons et al., 2016). Yet, the lower sensitivity of metabarcoding for rare species, potential biases from incomplete reference databases, and difficulties in distinguishing closely related taxa can limit its effectiveness for early warning purposes. This contrast in performance between approaches has been demonstrated across multiple empirical studies (Gargan et al., 2022; Wood et al., 2019a). For example, a comparison of qPCR and metabarcoding for detection of the invasive ascidian *Didemnum vexillum* Kott, 2002 (Gargan et al., 2022) showed that the targeted assay successfully detected the species in all eDNA samples from sites where it is known to occur in Ireland and Wales, whereas metabarcoding of the same samples failed to detect it, even at locations with established populations. This highlights the markedly higher sensitivity of targeted assays for early detection of priority invasives, while also confirming that metabarcoding remains valuable for capturing a broad range of native and non-native taxa in parallel. Similarly, in a comprehensive assessment of *S. spallanzanii*, qPCR and ddPCR consistently outperformed metabarcoding across both seawater and biofouling samples, with detection probabilities close to 1.0 for targeted assays compared to 0.27-0.57 for metabarcoding depending on the genetic marker used (Wood et al., 2019a). These findings reinforce a growing consensus: while metabarcoding offers unmatched breadth for community-wide biodiversity assessments, studies aiming to detect specific invasive or rare species, especially at low abundance or early stages of establishment, should prioritise targeted assays, or combine both approaches to balance sensitivity with taxonomic coverage.

### 3.4. Metazoans are the most frequently surveyed NIS via DNA

A total of 752 species has been identified using DNA-based methods across the 146 publications reviewed. The majority were metazoans (464 species), followed by Chromista (210 species) and plants (77), and a single species from Protozoa (**Fig. 4a**). Out of a total of 1,369 species detections reported across 146 publications (**Table S2**), approximately 31% represent new geographic records for the regions studied (**Table S2**; records on bold). This significant proportion highlights the value of DNA-based tools in enhancing the early detection of non-indigenous species in marine and coastal ecosystems. By revealing species that had not been previously recorded through traditional monitoring methods, these molecular approaches provide a powerful means to improve biosecurity and inform timely management interventions (Darling, 2019; Darling and Mahon, 2011; Duarte et al., 2021).

**Fig. 4.**
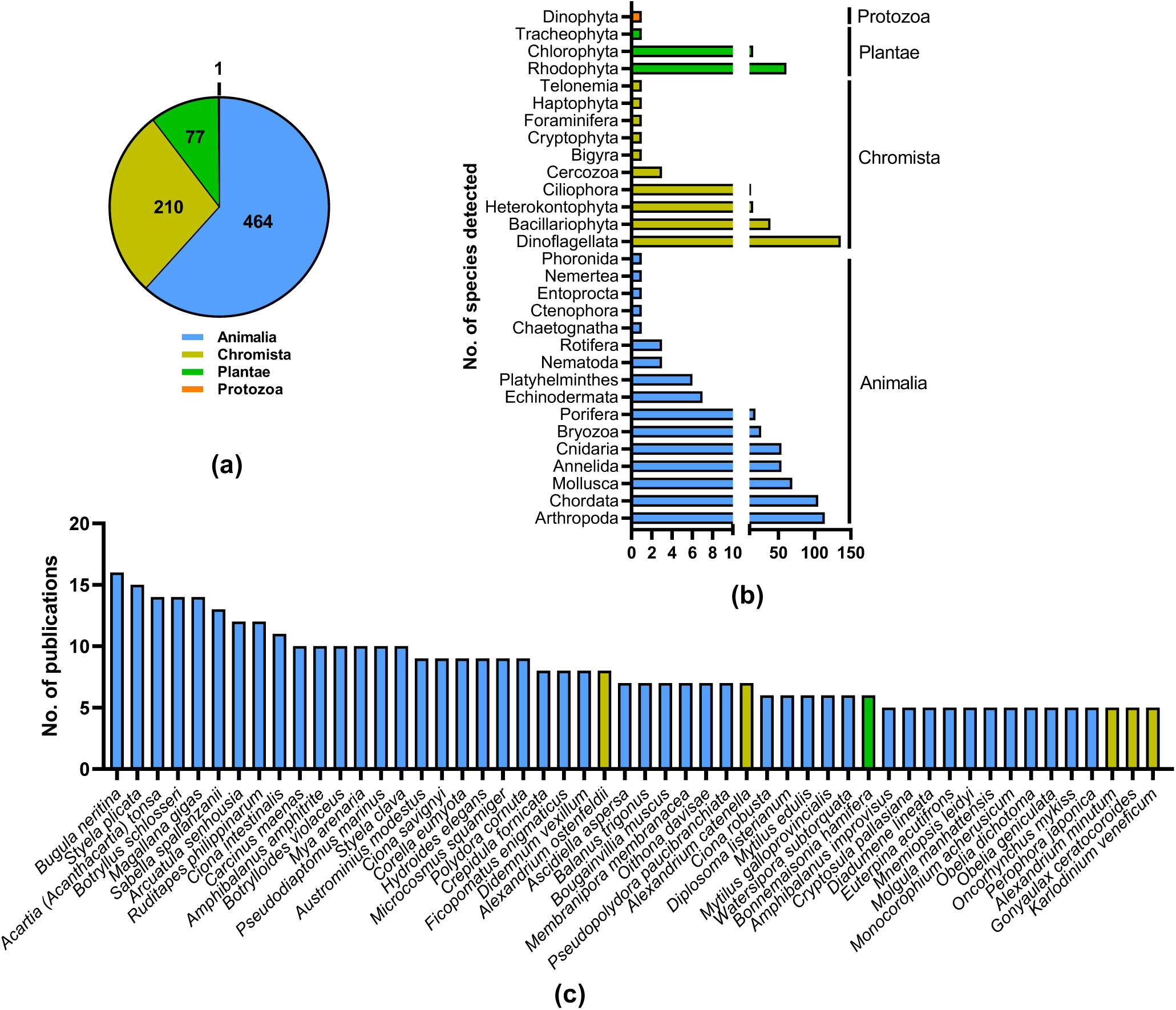
**(a)** Kingdoms and **(b)** phyla of all non-indigenous species detected through DNA-based tools in marine and coastal ecosystems and **(c)** most frequently detected species (at least in 5 publications).

Within Animalia, the most represented and ecologically relevant phyla were Arthropoda (**Fig. 4b**) (114 species), Chordata (105), and Mollusca (69) - groups that include numerous invasive taxa. Among Arthropoda, the classes Copepoda and Malacostraca (47 species each) were dominant (**Fig. S3**); both groups contain species known for their rapid dispersal, ecological plasticity, and role in ballast water transport (e.g., *Acarthia (Acanthartia) tonsa*, *Oithona davisae* Ferrari F.D. & Orsi, 1984, *Pseudodiaptomus marinus* Sato, 1913*, Monocorophium acherusicum* (A. Costa, 1853), *Grandidierella japonica* Stephensen, 1938, *Caprella scaura* Templeton, 1836, *Carcinus maenas* (Linnaeus, 1758)). In Chordata, the most represented classes were Teleostei (59 species), which includes several invasive fish (e.g., *Pterois miles* (Bennett, 1828), *Silurus glanis* Linnaeus, 1758, *Channa argus* (Cantor, 1842), *Cyprinus carpio* Linnaeus, 1758), and Ascidiacea (44), a class comprising many globally invasive tunicates (e.g., *Ciona intestinalis* (Linnaeus, 1767), *C. robusta* Hoshino & Tokioka, 1967, *S. plicata*, *Microcosmus squamiger* Michaelsen, 1927, *Didemnum vexillum* Kott, 2002). In Mollusca, the dominant classes were Bivalvia (43 species) and Gastropoda (22), both of which include taxa capable of altering benthic communities and fouling infrastructure (e.g., *Arcuatula senhousia* (W. H. Benson, 1842), *Crepidula fornicata* (Linnaeus, 1758), *Dreissena polymorpha* (Pallas, 1771), *Crepipatella dilatata* (Lamarck, 1822)).

Within Dinoflagellata, Dinophyceae (134 species) clearly dominated (**Fig. S3**), a group that includes several harmful algal bloom (HAB) species (e.g., *Alexandrium*, *Gymnodinium*) with significant ecological and economic consequences (Shang et al., 2019; Shaw et al., 2019; Sildever et al., 2022). Within Bacillariophyta (39), most species belonged to Bacillariophyceae (17) and Thalassiosirophyceae (15). These classes comprise ecologically important diatoms that contribute substantially to primary production and nutrient cycling, with some taxa (e.g. *Pseudo-nitzschia*) associated with harmful algal blooms, and others playing a key role in biofouling processes (Ardura et al., 2020; Sildever et al., 2022).

Within Plantae, most species belonged to Rhodophyta (61) and Chlorophyta (15) (**Fig. 4b**), with Florideophyceae (58 species) and Ulvophyceae (14) as the most represented classes, respectively (**Fig. S3**). These macroalgal groups include several invasive seaweeds known to displace native vegetation, alter habitats, and affect coastal food webs (e.g., *Codium fragile* (Suringar) Hariot 1889, *Caulerpa prolifera* (Forsskål) J.V.Lamouroux 1809*, Asparagopsis armata* Harvey 1855, *Bonnemaisonia hamifera* Hariot 1891) (Ibabe et al., 2024; Muha et al., 2019; Rey et al., 2020).

Most frequently reported non-indigenous species by DNA-based methods across the 146 publications (at least in 5 publications) were metazoans (**Fig. 4c**). Top-rank species were the bryozoan *B. neritina*, detected in 16 publications, and the sea squirt *S. plicata* (15 publications), both globally recognized marine biofoulers with a worldwide distribution. *Bugula neritina* has been recorded in harbours worldwide since at least 1921 (e.g., Pearl Harbor) (Carlton and Eldredge, n.d.; Coles et al., 1999) and exhibits strong tolerance to copper-based antifouling agents, enabling it to thrive on ship hulls, aquaculture gear, and artificial structures. *Styela plicata*, known for its high abundance on submerged infrastructures in temperate to tropical waters, has been widely documented across North America, the Mediterranean, and Asia, where it outcompetes native fauna for space and fouls cultured oysters and mussels (Mckindsey et al., 2007).

Several factors likely contribute to the frequent detection of these species in DNA-based studies. First, their high abundance in biofouling communities associated with marinas, ports, and aquaculture facilities, common sampling sites in eDNA and metabarcoding surveys, increases the probability of detection (Ammon et al., 2018; Andrés et al., 2023; Ardura et al., 2021a; Aylagas et al., 2024; Ibabe et al., 2021; Kim et al., 2018; Lavrador et al., 2024; Obst et al., 2020; Pearman et al., 2021; Rey et al., 2020; Zarcero et al., 2024) (**Table S2**). Second, both species are well represented in public genetic reference databases, which improves taxonomic assignment efficiency. For instance, *B. neritina* has 695 sequences available, including 208 COI and eight 18S entries, on the Barcode of Life DataSystem (https://v4.boldsystems.org/; accessed on 28^th^ January 2026), while *S. plicata* is represented by 190 sequences, with 172 COI and eight 18S entries. This strong coverage of commonly used barcoding markers facilitates DNA-based identification. Third, their sessile or semi-sessile nature results in persistent DNA signals in the environment, making them especially suitable for eDNA detection (Couton et al., 2022). Other commonly reported species include the copepod *A. tonsa* (14 publications) (Andrés et al., 2023; Lavoie et al., 1999; Ware et al., 2016), known for rapid dispersal in ballast water; the colonial tunicate *B. schlosseri*, noted for its invasive spread and fouling behaviour (14 publications) (Sheets et al., 2016; Zwahlen et al., 2022); and the bivalve *Magallana gigas* (Thunberg, 1793) (14 publications), an aquaculture species notorious for its ecological impacts due to habitat alteration and competition (Des et al., 2022; Hansen et al., 2023). *Sabella spallanzanii* appear as fourth in the list (**Fig. 4c**) (13 publications) (Ammon et al., 2018; Brand et al., 2022; Wood et al., 2019b, 2019a), likely due to its status as a well-known invasive polychaete and a frequent target of DNA-based monitoring. The species has been the focus of several specific detection studies, particularly in Oceania, most notably in New Zealand, where it is a major biosecurity concern (Ammon et al., 2018; Scriver et al., 2024; Wood et al., 2019b, 2019a).

### 3.5. Water is the most used sample type for DNA-based NIS detection

Water has been the most adopted sample type for surveying non-indigenous species in marine and coastal ecosystems, with over 60% of the reviewed publications reporting its use (**Fig. 5a, Table S2**) (Ardura et al., 2020; Borrell et al., 2018; Couton et al., 2022; Ellis et al., 2022; Holman et al., 2019; Suarez-Menendez et al., 2020; Westfall et al., 2020; Wood et al., 2019b). Of these, about 77% reported its exclusive use (33% in metabarcoding and 44% for single species detection) (Ellis et al., 2022; Fernandez et al., 2022; Gargan et al., 2022; Maggio et al., 2023; Suarez-Menendez et al., 2020). This prevalence is not unexpected due to several reasons: 1) water sampling is relatively simple, non-destructive, and requires minimal specialized equipment. It can be quickly collected using basic containers, Niskin bottles, or peristaltic pumps, making it feasible even in remote or challenging environments (Lacoursière-Roussel et al., 2018) or onboard (Ardura et al., 2020, 2015a). Environmental DNA in water integrates genetic material shed by organisms through skin, scales, mucus, feces, gametes, and decomposing tissues (Deiner et al., 2017). This allows detection of a wide range of taxa - from microbes to large vertebrates – and, thus, the early detection of NIS belonging to very different taxonomic groups (e.g., metazoans, microalgae, microeukaryotes) - without direct observation or capture (Ardura et al., 2021b). Water sampling has become widely adopted in eDNA workflows, leading to protocols that are more established and reproducible for filtration, preservation, and DNA extraction, which enhances comparability across studies (Goldberg et al., 2016, 2015; Ruppert et al., 2019).

**Fig. 5.**
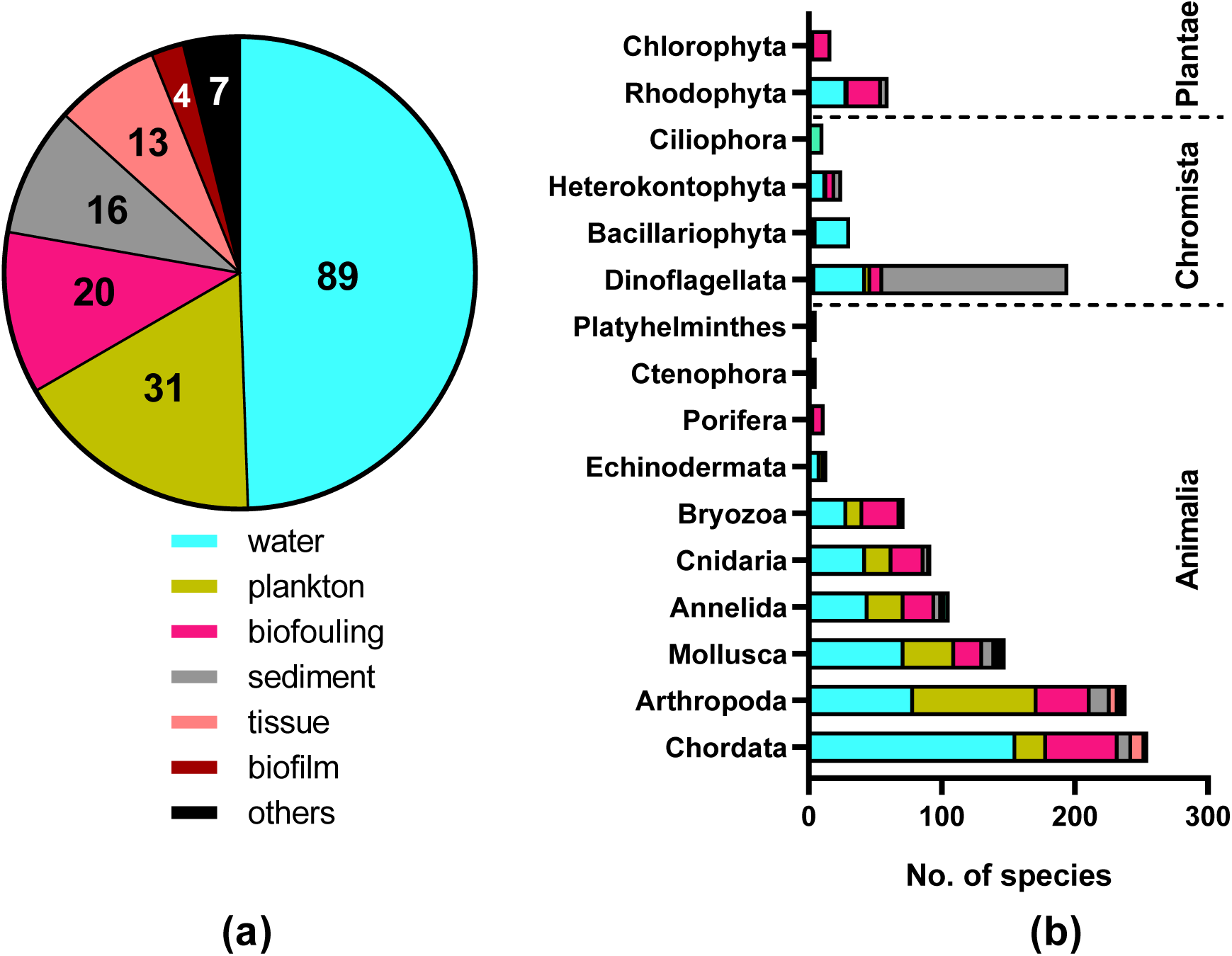
**(a)** Types of samples or substrates collected for the detection of non-indigenous species using DNA-based tools, with the number of publications for each sample type; and **(b)** number of species detected per phylum for each sample type.

The second most employed substrate is zooplankton (**Fig. 5a**), which may also offer several practical and ecological advantages. Many marine NIS are introduced during their planktonic larval stages (in particular Arthropoda-Copepoda), which are abundant in zooplankton samples and often morphologically indistinct, making molecular identification especially valuable for early detection (Abad et al., 2016; Berry et al., 2023; Brown et al., 2016; Rey et al., 2020; Zaiko et al., 2015). Zooplankton is easy to collect with simple gear like plankton nets, allowing broad spatial and temporal coverage, especially in high-risk areas like ports and marinas (Brown et al., 2016; Darling et al., 2018; Lin et al., 2022). These samples usually contain sufficient biomass for efficient DNA extraction and amplification, and their mixed nature is well-suited for DNA metabarcoding, enabling the simultaneous detection of multiple taxa, including rare or cryptic species (Abad et al., 2016; Brown et al., 2016; Darling et al., 2018; Lin et al., 2022) or single species detection (Richardson et al., 2016; Simpson et al., 2023; Uthicke et al., 2018; Wood et al., 2017). Additionally, zooplankton communities can reflect rapid changes in biodiversity, making them ideal for monitoring community shifts due to biological invasions (Abad et al., 2016).

Less commonly used sample types include biofouling and sediment (**Fig. 5a**), which often require specialized methods such as divers, dredges, or settlement plates; the latter needing an additional colonization period of several months. These factors make such approaches less scalable and cost-efficient for large-scale monitoring programs (Altermatt et al., 2023). However, despite these challenges, biofouling and sediment samples can be particularly valuable for detecting NIS because they target organisms that settle and accumulate on hard substrates or within sediments, including early life stages and cryptic species many of which may be missed in water-based eDNA or zooplankton samples (Antich et al., 2020). These sample types can therefore enhance detection sensitivity and provide complementary information on benthic and sessile NIS communities, improving overall monitoring effectiveness (Koziol et al., 2019; Lavrador et al., 2024; Rey et al., 2020). Indeed, a detailed analysis of the most dominant taxonomic groups detected in each sample type revealed considerable differences: while Chordata have been mostly detected in water samples, Arthropoda are preferentially detected in zooplankton samples, Dinoflagellata are by far the most abundant in sediments, and Rhodophyta dominate fouling communities, reflecting the distinct ecological niches and life histories captured by each sampling method (**Fig. 5b**). This suggests that employing a multi-substrate sampling approach may improve the overall detection of non-indigenous species (NIS) by capturing a broader range of taxa (Fernandez et al., 2021; Koziol et al., 2019; Lavrador et al., 2024; Rey et al., 2020; Ríos-Castro et al., 2021). Nevertheless, relatively few studies have implemented such integrative strategies, likely due to logistical and financial constraints (82% of the publications used one sample type; 14% 2 sample types and 2% used 3 and 4 sample types). A Venn diagram illustrating the partitioning of species among the most used substrates - water, sediment, plankton, and biofouling/film - further supports this observation: only about 3% of species were detected across all four substrates. In contrast, water samples exclusively accounted for 31% of species detections, biofouling/film for 18%, plankton for 11%, and sediment for 15%, highlighting the complementary nature of these sample types and the limitations of relying on a single substrate for comprehensive NIS surveillance (**Fig. S4**).

### 3.6. Marker region usage in DNA-based NIS surveys

The mitochondrial cytochrome c oxidase subunit I (COI) has been the most adopted marker for surveying non-indigenous species in marine and coastal ecosystems (73% of the reviewed publications; **Fig. 6a, Table S2**) (Borrell et al., 2018; Fernandez et al., 2021; Kim et al., 2018; Lavrador et al., 2024; Wood et al., 2017; Zarcero et al., 2024). This widespread use is largely due to COI broad applicability across metazoans, particularly animals such as chordates, crustaceans, and molluscs, which constitute the majority of surveyed NIS (**Fig. 4** and **Fig. 6b**). Its high species-level resolution, combined with extensive representation in reference databases (e.g., BOLD, GenBank) (Duarte et al., 2020; Weigand et al., 2019), allows for accurate taxonomic assignment. Furthermore, the mitochondrial origin of COI ensures high copy numbers per cell, increasing detection sensitivity in both bulk tissue and environmental DNA samples. The availability of universal primers (**Table S3**) and standardized protocols facilitates consistent and comparable monitoring across regions and time, which is essential for early detection and management of invasive species. Within COI, and on metabarcoding studies, the primer pair mlCOIintF/jgHCO2198 (Geller et al., 2013; Leray et al., 2013), targeting a short fragment (∼313 bp), has become by far the most frequently employed in metabarcoding studies aiming to detect non-indigenous species (NIS) in marine and coastal ecosystems (31 publications, **Table S3**). This predominance results from its broad metazoan coverage, high amplification efficiency, and the relatively short amplicon length that performs well with degraded environmental DNA while retaining sufficient taxonomic resolution for species-level identification (Geller et al., 2013; Leray et al., 2013).

**Fig. 6.**
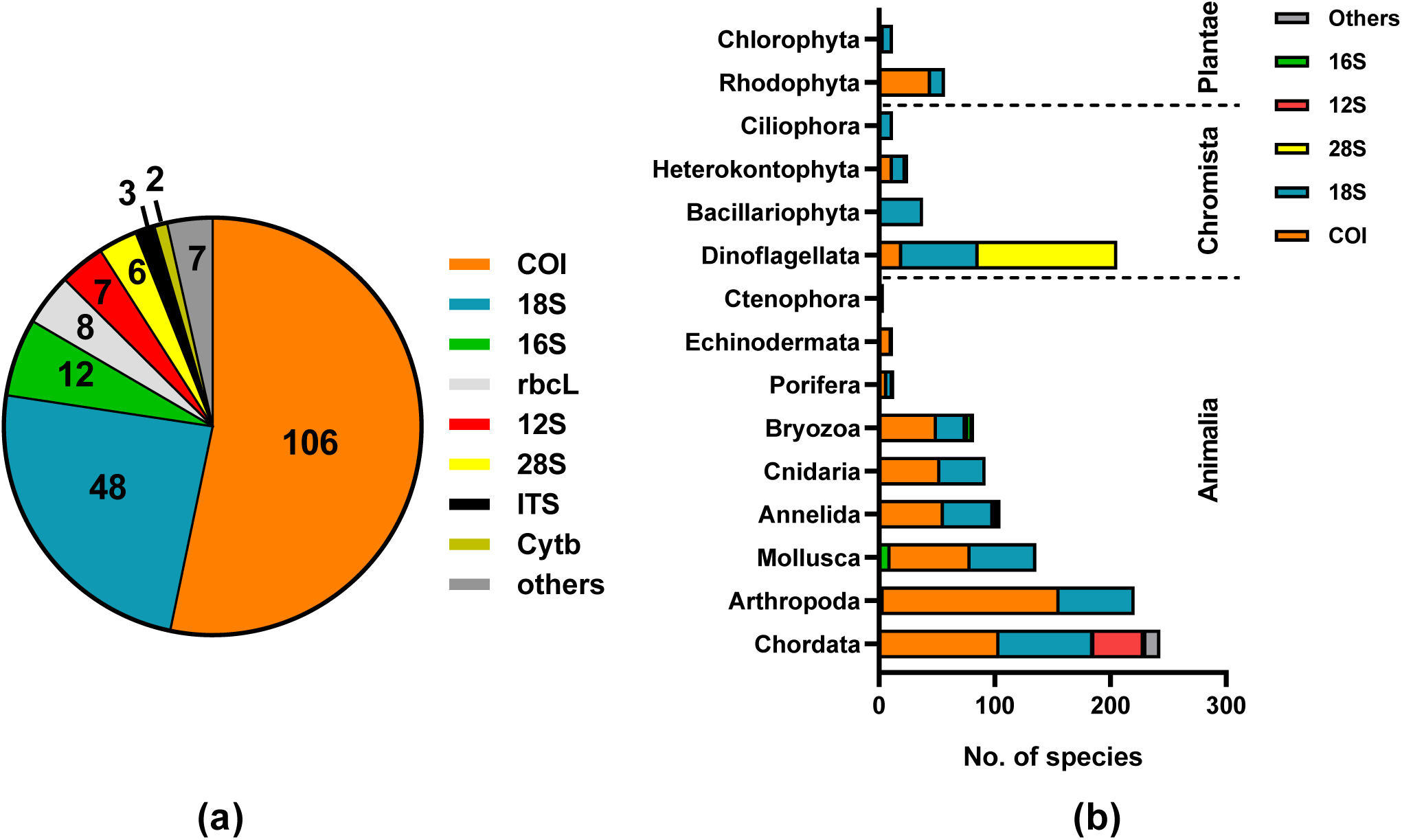
**(a)** Genetic markers used for the detection of non-indigenous species using DNA-based tools, with the number of publications for each marker and **(b)** number of species detected per phylum for each marker.

Other widely used genetic markers include 18S rRNA (33% of the publications) (Abad et al., 2016; Ammon et al., 2018; Brown et al., 2016; Darling et al., 2018; Holman et al., 2019) and mitochondrial 16S rRNA (8% of the publications) genetic markers (Ardura et al., 2017; Berry et al., 2023; Wang et al., 2024). The 18S gene is commonly used for broad eukaryotic surveys, while the mitochondrial 16S marker has been particularly valuable as an alternative or complement to COI-based approaches for metazoan detection (Berry et al., 2023; Couton et al., 2022; Koziol et al., 2019; Westfall et al., 2020). Within 18S, the Uni18SF/Uni18SR (V4) (16 publications) (Zhan et al., 2013) and SSU_F04/SSU_R22 (V1–V2) primer pairs (6 publications) (Blaxter et al., 1998) (**Table S3**), have also been widely used for the detection of a broader range of eukaryotes, including non-metazoan taxa. However, despite its broad taxonomic coverage, the 18S marker generally provides limited species-level resolution due to its conserved nature (Karst et al., 2018; Wu et al., 2015), which can hinder accurate discrimination among closely related taxa - an important limitation in the context of non-indigenous species monitoring. Although the 18S-V9 region has been described as having higher taxonomic resolution and the potential to detect a greater diversity of organisms, its limited reference coverage compared to the V4 region currently constrains its ability to provide detailed taxonomic annotations (3 primer pairs used in a total 6 publications, **Table S3**) (Choi and Park, 2020).

In addition, several studies have targeted group-specific markers to improve detection efficiency and taxonomic resolution within taxa. For example, the 12S rRNA gene, in particular the MiFish primers (Miya et al., 2015), has been extensively used for the detection and monitoring of fish species (Cananzi et al., 2022; Jannel et al., 2025; Spence and Skelton, 2024), while the 28S rRNA gene has been applied for the monitoring of harmful algal species (Dias et al., 2015; Shang et al., 2024, 2019) (**Fig. 6b**).

In the scope of metabarcoding, the use of multiple genetic markers can substantially increase the taxonomic breadth of detected species by capturing organisms that may be missed by a single locus (Duarte et al., 2023b; Lavrador et al., 2024; Westfall et al., 2020). However, most studies to date have relied on a single genetic marker (>70% of the publications), whereas a smaller proportion employed two markers (19%), and only 7% used three or more within the same study. A Venn-diagram illustrating the overlap among the most frequently used markers - COI, 28S, 18S, 16S, and 12S - showed that no species were detected by all five markers (**Fig. S5**). In contrast, COI exclusively accounted for 35% of species detections, 18S for 28%, 28S for 11%, 12S for 5%, and 16S for 0.5%, highlighting both the complementary nature of genetic markers and the limitations of relying on a single locus for comprehensive NIS surveillance. It is important to note that this partitioning is based on different publications, each of which may have selected the marker best suited to the target communities (and thus may explain why no species have been detected by all markers). Thus, this lack of complete overlap is not unexpected, given the differing taxonomic resolution and target groups of each marker. For instance, 12S is primarily used for vertebrate (especially fish) detection (Kume et al., 2021; Miya et al., 2015; Spence and Skelton, 2024), and therefore its limited ability to detect invertebrates is inherent to its design. In contrast, markers such as COI and 18S are more broadly applied to invertebrate communities (Duarte et al., 2023b; Grey et al., 2018; Lavrador et al., 2024; Rey et al., 2020). Additionally, 16S, although mitochondrial in animals, can vary in taxonomic coverage depending on primer choice and reference database completeness, further contributing to differences in species detection across markers (Elbrecht et al., 2016). Nevertheless, even in studies that employ multiple markers the overlap of detected species is very low (Castro-Cubillos et al., 2022; Couton et al., 2022; Moutinho et al., 2023; Rey et al., 2020; Simões et al., 2024; Von Ammon et al., 2023; Westfall et al., 2020), further supporting that a multi-marker metabarcoding strategy is the most appropriate for broad-spectrum monitoring and minimizing false negatives, which are particularly critical when surveying NIS. Therefore, while COI remains the most informative marker for species-level identification of metazoan NIS, 18S and 16S markers continue to play an important complementary role, particularly for assessing overall eukaryotic community composition.

Across this literature review, a total of 78 species-specific assays targeting 65 non-indigenous species were identified (**Fig. 7; Table S4**), indicating that multiple assays have been developed for some high-priority taxa. Most species targeted on these assays were metazoans (79%), Chromista (15%) and Plantae (6%) (**Fig. 7a**), reflecting both the prevalence of animal invaders in aquatic ecosystems and the stronger historical focus of molecular monitoring efforts on metazoans. These assays covered a broad taxonomic range (**Fig. 7b**), including taxa of high ecological and socio-economic relevance such as fish (Knudsen et al., 2022), bivalves (Ardura et al., 2017; Yip et al., 2021), crustaceans (Knudsen et al., 2022) or macroalgae (Knudsen et al., 2022). Several assays targeted globally recognised high-risk invaders, including *C. maenas* (Crane et al., 2021; Danziger and Frederich, 2022), *C. carpio* (Takahara et al., 2012), *D. polymorpha* (Ardura et al., 2017), *Asterias amurensis* Lutken, 1871 (Yang et al., 2024), *Eriocheir sinensis* H. Milne Edwards, 1853 (Knudsen et al., 2022), *Mnemiopsis leidyi* A. Agassiz, 1865 (Knudsen et al., 2022), *Oncorhynchus mykiss* (Walbaum, 1792) (Fu’adil Amin et al., 2021; Knudsen et al., 2022), *Undaria pinnatifida* (Harvey) Suringar, 1873 (Bott et al., 2015) and *S. spallanzanii* (Wood et al., 2017), among others, listed among the 100 of the World’s Worst Invasive Alien Species by the IUCN / ISSG (https://portals.iucn.org/library/sites/library/files/documents/2000-126.pdf). The frequent targeting of these species reflects their wide geographic spread, severe ecological impacts, and the need for early detection tools to support management and control strategies (Bott et al., 2015). Most assays targeted COI region (65%), consistent with its high interspecific variability, extensive reference databases, and widespread use in DNA barcoding of animal species, as already explained above. Other loci were used less frequently for developing specific assays (<8% each), including 28S rRNA, CytB, 16S rRNA, ITS and rbcL, reflecting taxon-specific constraints and the need to balance assay specificity with cross-taxon primer compatibility (**Fig. 7c**).

**Fig. 7.**
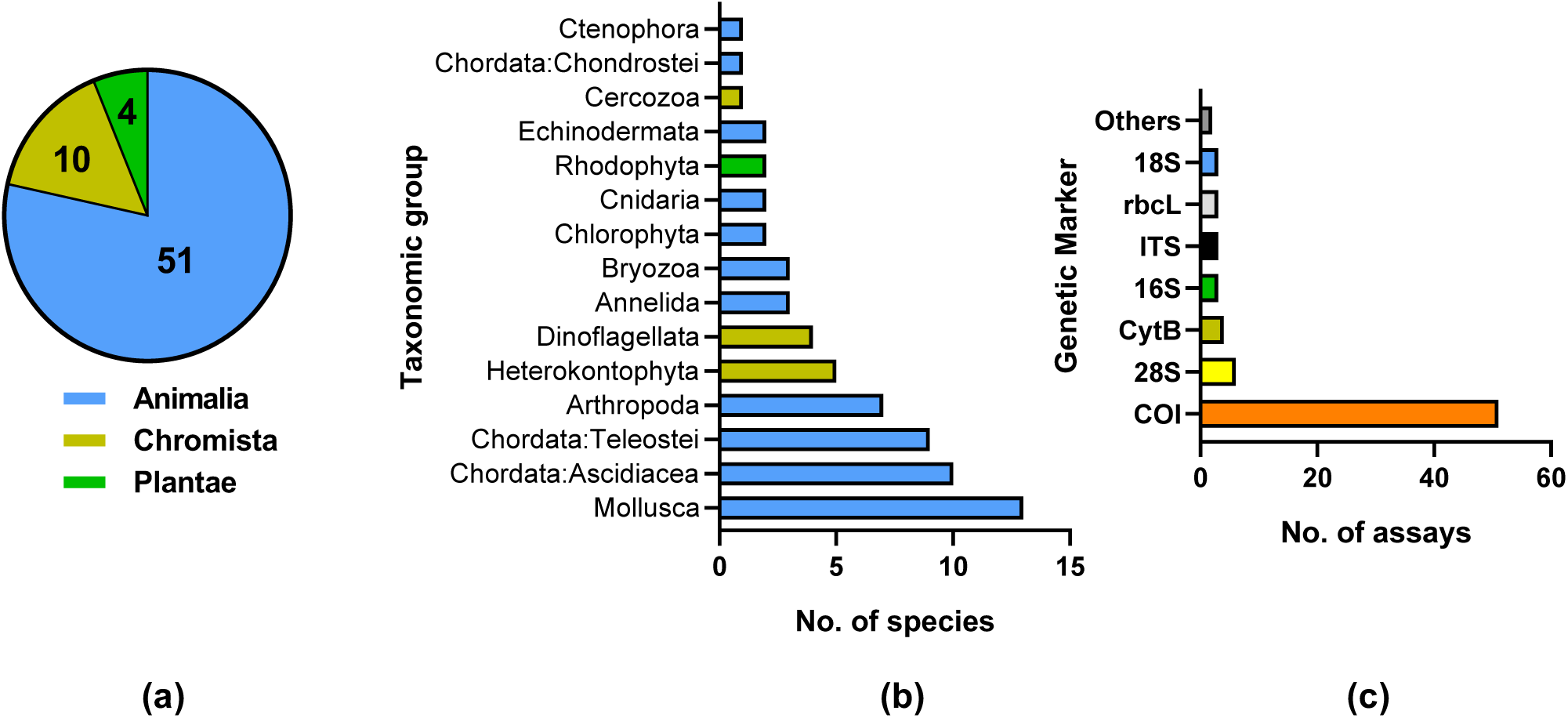
**(a)** Kingdoms and **(b)** phyla for which DNA-based, species-specific assays have been developed for the detection of non-indigenous species, and **(c)** the genetic markers most used in these assays.

Amplicon lengths span from very short fragments (<100 bp) optimised for degraded eDNA, to longer targets (>300 bp) in assays developed for tissue-derived DNA or early eDNA applications with conventional PCR (Ardura et al., 2017, 2015b; Vercaemer et al., 2015; Willis et al., 2011) (**Table S4**). Most assays were developed for qPCR, which remains the predominant approach due to its accessibility (**Fig. 3**), high sensitivity, and ability to provide quantitative or semi-quantitative estimates of target DNA. More recent studies have increasingly adopted droplet digital PCR (ddPCR), taking advantage of its enhanced analytical sensitivity, absolute quantification, and reduced susceptibility to PCR inhibition; features that are particularly valuable when working with turbid waters or complex environmental matrices (Brand et al., 2022; Crane et al., 2021; Uthicke et al., 2018; Wood et al., 2019a).

## 5. Implications for marine biosecurity and monitoring programmes

The findings of this global synthesis have direct implications for the design and implementation of marine biosecurity and invasive species monitoring programmes (**Fig. 8**). First, the high proportion of new regional records detected using DNA highlights its value as an early warning tool capable of identifying incursions before populations become established. Incorporating DNA-based approaches into routine surveillance can therefore substantially improve prevention and rapid response strategies, which are widely recognised as the most cost-effective stages for invasive species management (Morisette et al., 2021).

**Fig. 8.**
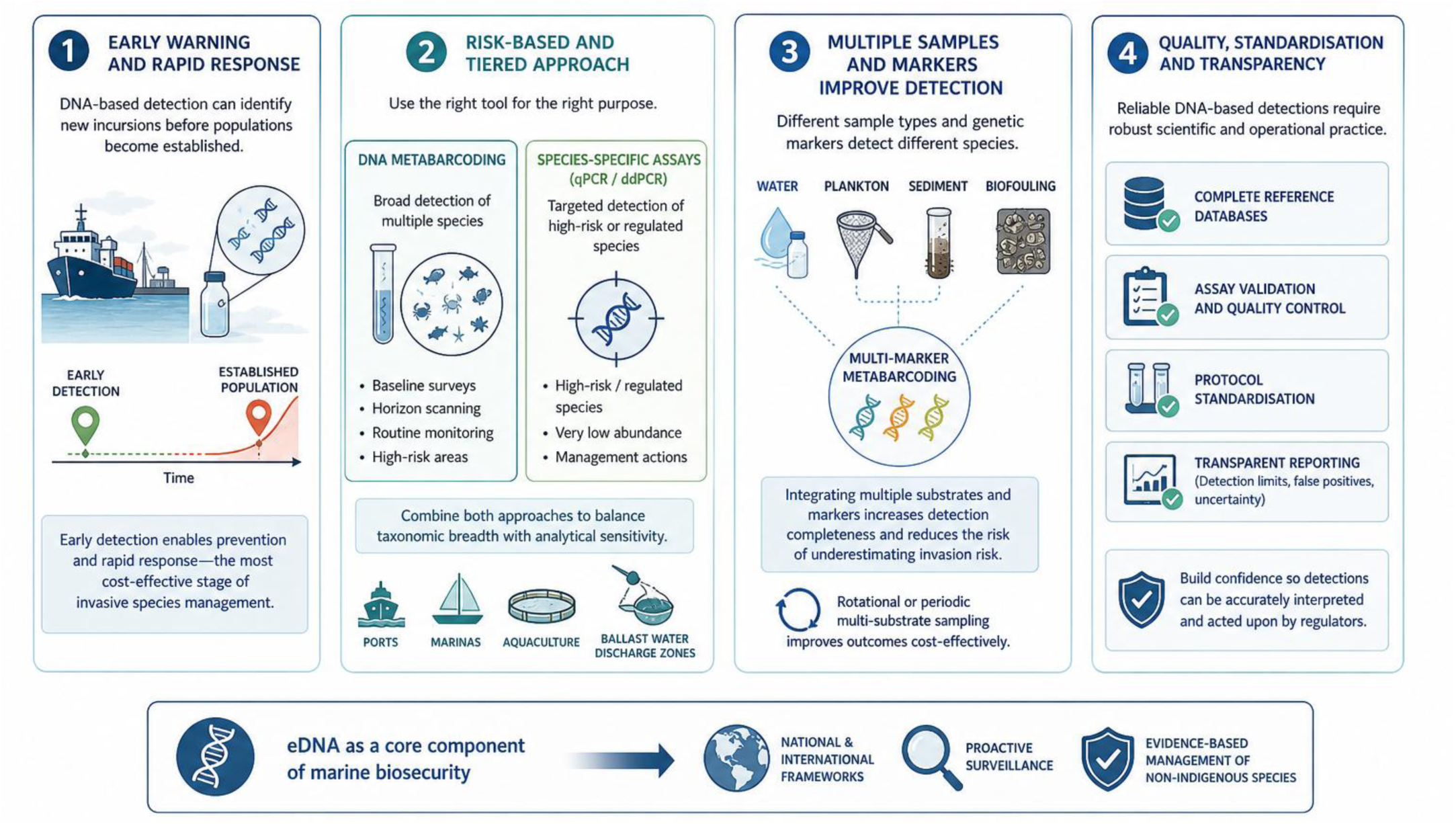
Conceptual framework for integrating DNA-based tools into marine biosecurity and invasive species monitoring. Figure created using an AI-based image generation tool (ChatGPT, OpenAI), based on authors-defined scientific content.

Second, current results indicate that effective biosecurity monitoring should adopt a risk-based and tiered approach. DNA metabarcoding is well suited for baseline surveys, horizon scanning, and routine monitoring in high-risk areas such as ports, marinas, aquaculture facilities, and ballast water discharge zones, where propagule pressure is highest (Ammon et al., 2018). In contrast, species-specific assays (qPCR or ddPCR) should be prioritised for targeted surveillance of high-risk or regulated species, particularly when management actions depend on confident detection at very low abundance (Duarte et al., 2023a; Wood et al., 2019b). Thus metabarcoding approaches are potentially useful at detecting multiple NIS in a single analysis and are therefore useful for early detection and as supplementary survey tools. Combining both approaches within the same monitoring framework allows managers to balance taxonomic breadth with analytical sensitivity (Gargan et al., 2022; Simmons et al., 2016).

Third, the strong complementarity observed among sample types and genetic markers suggests that reliance on single-substrate or single-marker strategies may underestimate invasion risk (Koziol et al., 2019; Lavrador et al., 2024; Rey et al., 2020; Zarcero et al., 2024). Where resources permit, integrating water sampling with plankton, sediment, or biofouling surveys, alongside multi-marker metabarcoding, can markedly improve detection completeness. Even limited rotational or periodic multi-substrate sampling could substantially enhance surveillance outcomes without prohibitive cost increases (Lavrador et al., 2024).

Finally, the operational employment of DNA-based approaches in biosecurity contexts requires continued investment in reference database completeness, assay validation, and protocol standardisation. Transparent reporting of detection limits, false positives, and uncertainty is essential to ensure that DNA-based detections can be confidently interpreted and acted upon by regulatory agencies (Darling, 2019; Mosher et al., 2020; Sepulveda et al., 2020). As these challenges are progressively addressed, eDNA is well positioned to become a core component of national and international marine biosecurity frameworks, supporting proactive, evidence-based management of non-indigenous species.

## 6. Conclusions

This global synthesis demonstrates that DNA-based approaches are now a powerful and operationally relevant component of non-indigenous species surveillance in marine and coastal ecosystems. Across 146 studies, DNA-based tools enabled the detection of 752 non-indigenous species spanning multiple taxonomic groups, with nearly one third representing new regional records. This highlights the strong value of DNA for early warning and biosecurity, particularly where traditional monitoring is limited by logistics or taxonomic constraints. Research effort remains geographically uneven, with a strong concentration in Europe, Oceania and North America, and major gaps across regions such as Africa, the Arctic, and Antarctica. From a biosecurity perspective, expanding DNA-based monitoring into these under-surveyed areas should be a priority, given increasing maritime traffic and coastal development. Methodologically, capturing the full spectrum of marine invasive taxa remains challenging. Species-specific assays provide higher sensitivity for early detection of a small set of priority invaders, while DNA metabarcoding enables broad surveillance and the exposure of untargeted species. However, minimalistic approaches - such as relying on a single marker, substrate, or limited sampling effort - although logistically attractive, are often insufficiently rigorous and can substantially increase the risk of false negatives. This limitation is particularly critical in the context of non-indigenous species, where failure to detect an incipient population may compromise timely management action. Consequently, multi-substrate and multi-marker strategies consistently improve detection completeness compared to simplified designs, yet remain underused. Importantly, shifting the focus from maximising sampling coverage to optimising detection reliability is essential; more integrative and methodologically robust approaches applied to fewer locations may yield more informative and reliable results than extensive but analytically limited surveys. Overall, DNA-based approaches have clearly moved beyond proof-of-concept and is increasingly supporting real-world biosecurity and management applications. Continued investment in integrated monitoring frameworks, reference database development, protocol standardisation, and coordinated international surveillance will be essential to fully realise the potential of DNA for the early detection and management of marine biological invasions.

## 7. Ethical Statement

This review did not involve the collection of new data from humans or animals and therefore did not require ethical approval.

## Author Contributions

Sofia Duarte: Conceptualization; Data curation; Formal analysis; Funding acquisition; Investigation; Methodology; Project administration; Resources; Software; Supervision; Validation; Visualization; Roles/Writing - original draft; Writing - review & editing. Filipe O. Costa: Writing- Reviewing and Editing.

## Declaration of generative AI and AI-assisted technologies in the manuscript preparation process

During the preparation of this work, the authors used ChatGPT (OpenAI) to assist with language editing, improving the clarity, grammar, and readability of the manuscript, as well as to support the creation of the graphical abstract and Figure 8 based on author-defined scientific content. No scientific content, data interpretation, or conclusions were generated using this tool. All outputs were carefully reviewed and edited by the authors, who take full responsibility for the content of the published article.

