## Supplementary material 1 for "Tracing the intruders: a global appraisal of marine invasive species detection through DNA-based approaches"

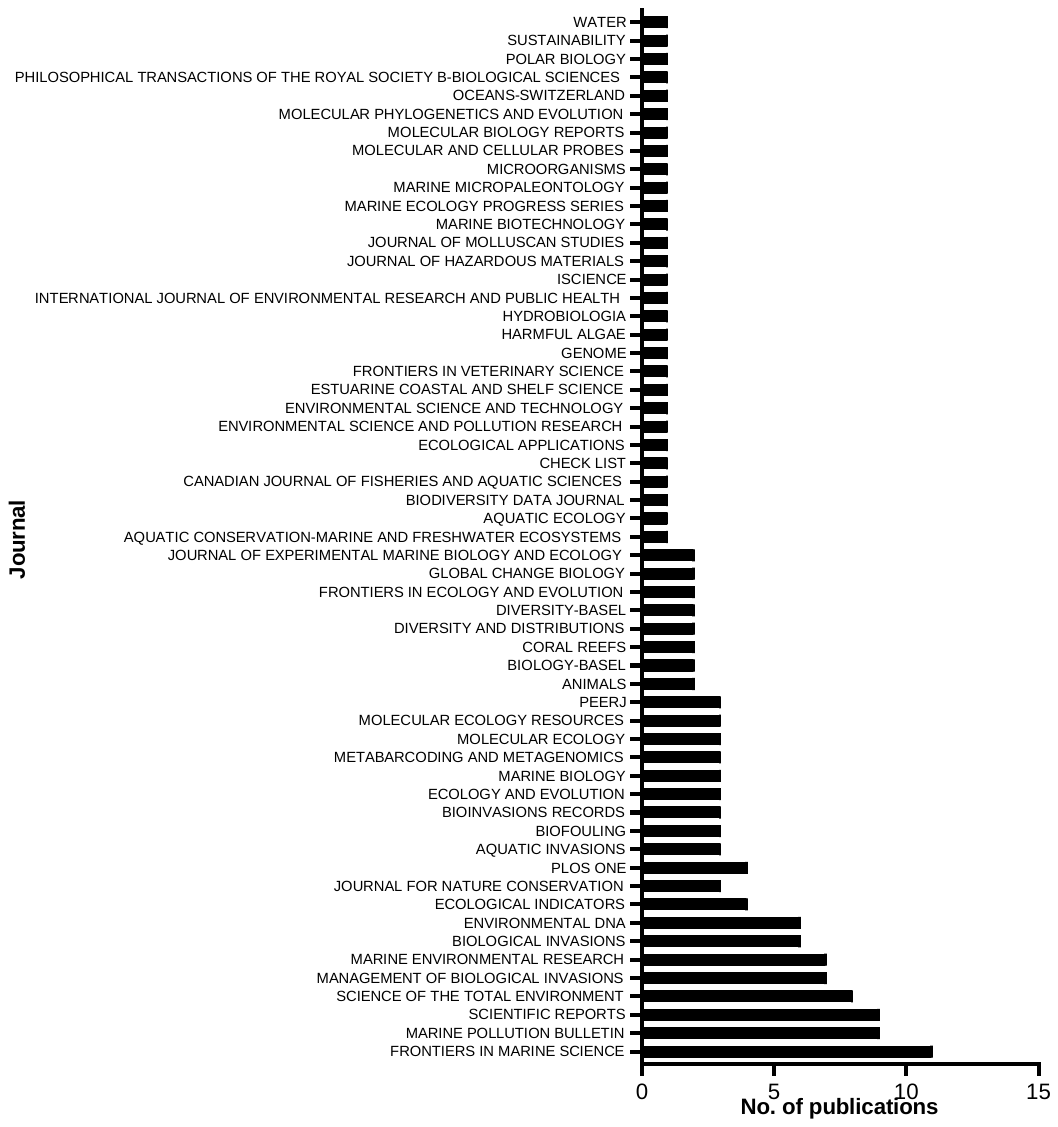

**Fig. S1.** Number of publications per journal featuring the 146 experimental studies on the use of DNA-based tools for the detection and monitoring of non-indigenous species in marine and coastal ecosystems.

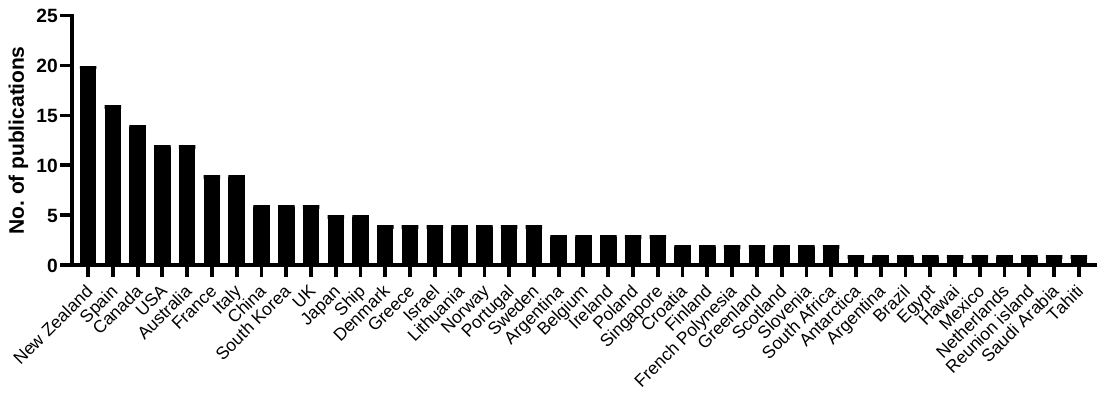

**Fig. S2.** Number of publications per country surveyed using DNA-based tools for the detection and monitoring of non-indigenous species in marine and coastal ecosystems.

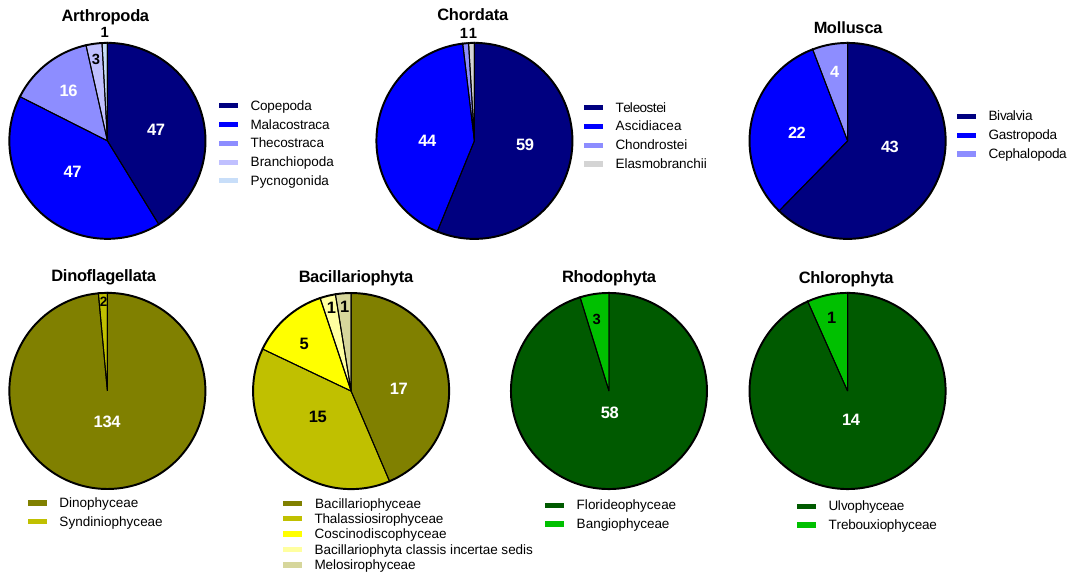

**Fig. S3.** Most represented classes within the dominant phyla of each kingdom.

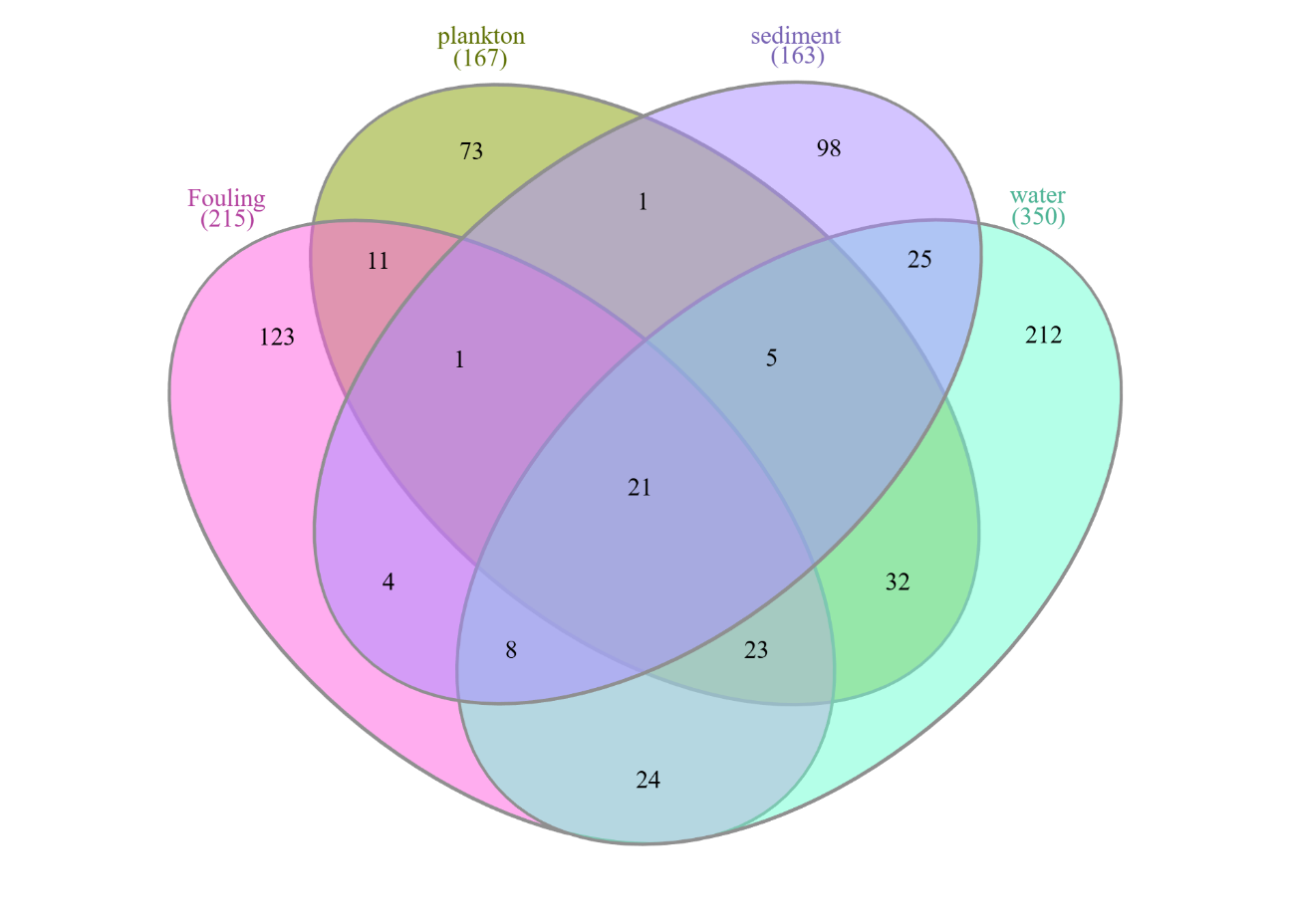

**Fig. S4.** Partitioning of the marine non-indigenous species (NIS) detected in the most used sample types (water, biofouling/film, plankton and sediment). For this analysis, only publications that clearly specified which species were detected on each substrate were considered.

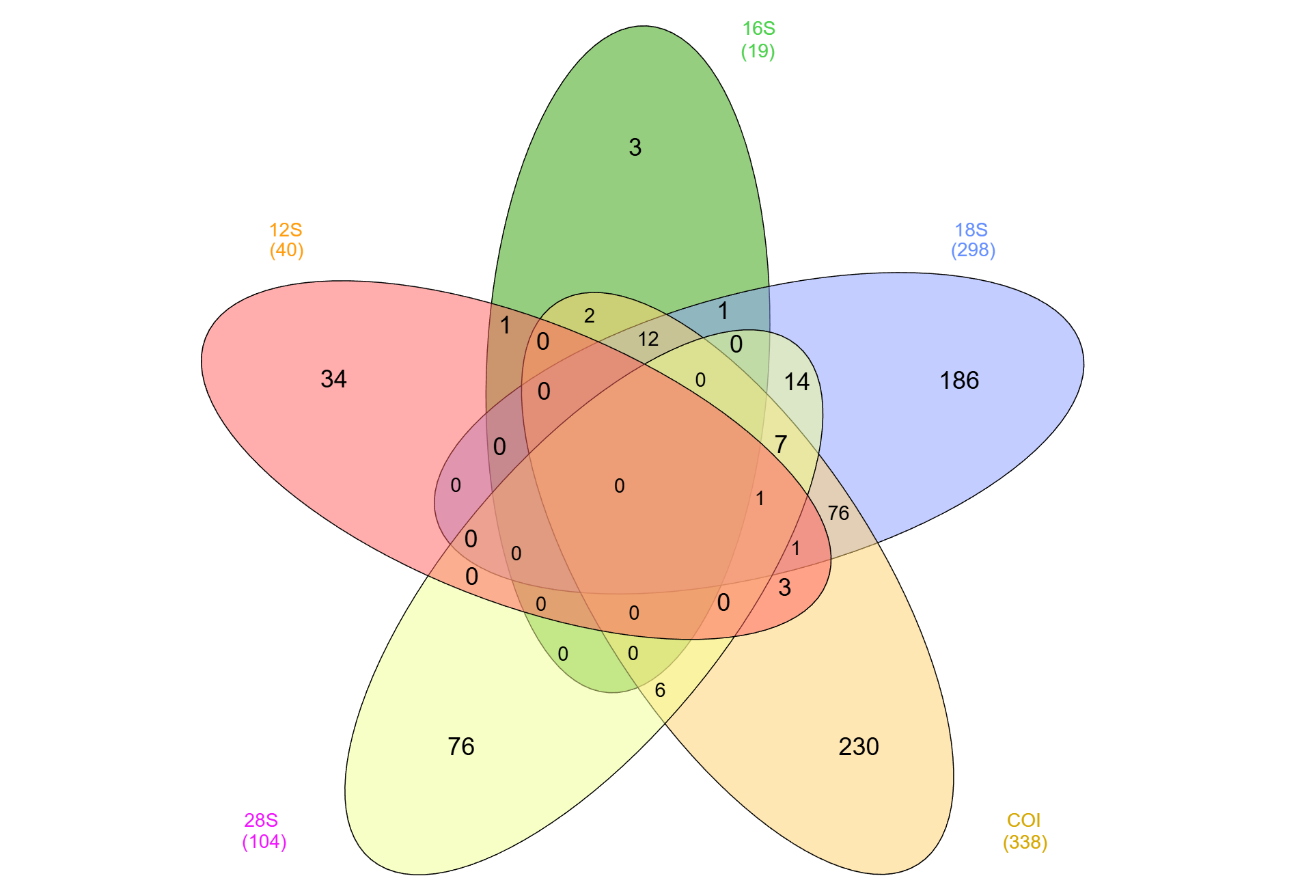

**Fig. S5.** Partitioning of the marine non-indigenous species (NIS) detected using the most employed genetic markers (COI, 28S, 18S, 12S and 16S). For this analysis, only publications that clearly specified which species were detected using each marker were considered.
